# BUMBLE BEE RESPONSES TO CLIMATE AND LANDSCAPES: INVESTIGATING HABITAT ASSOCIATIONS AND SPECIES ASSEMBLAGES ACROSS GEOGRAPHIC REGIONS IN THE UNITED STATES OF AMERICA

**DOI:** 10.1101/2023.10.16.562230

**Authors:** Morgan E. Christman, Lori R. Spears, Emily K. Burchfield, William D. Pearse, James P. Strange, Ricardo A. Ramirez

## Abstract

Bumble bees are integral pollinators of native and cultivated plant communities, but species are undergoing significant changes in range and abundance on a global scale. Climate change and land cover alteration are key drivers in pollinator declines; however, limited research has evaluated the cumulative effects of these factors on bumble bee assemblages. This study tests bumble bee responses to climate and land use by modeling 1) occupancy (presence/absence); 2) species richness; 3) species-specific habitat requirements; and 4) assemblage-level responses across geographic regions. We integrated richness, abundance, and distribution data for 18 bumble bee species with site specific bioclimatic, landscape composition, and landscape configuration data to evaluate the effects of multiple environmental stressors on bumble bee assemblages throughout 433 agricultural fields in Florida, Indiana, Kansas, Kentucky, Maryland, South Carolina, Utah, Virginia, and West Virginia from 2018 to 2020. Increased prevalence of attractive crops was associated with increased bumble bee presence, while higher maximum temperature of warmest month and unattractive crops were linked to bumble bee absences. Bumble bee species richness was positively correlated with attractive crops and elevation but declined with precipitation of the wettest month. Moreover, species richness increased as maximum temperature of warmest month approached 29°C but declined at they rose to 36°C, suggesting a potential temperature threshold around 33°C. Distinct east vs. west groupings emerged when evaluating species-specific habitat associations, prompting a detailed evaluation of bumble bee assemblages by geographic region. Overall, climate and land use combine to drive bumble bee occupancy and assemblages, but how those processes operate is idiosyncratic and spatially contingent across regions. From these findings, we suggested regionally specific management practices to best support rich bumble bee assemblages in agroecosystems. Results from this study contribute to a better understanding of climate and landscape factors affecting bumble bees and their habitats throughout the USA.

## 1. INTRODUCTION

Bumble bees (Hymenoptera: Apidae: *Bombus* Latreille) are essential, globally widespread pollinators of native and cultivated plant communities (Kremen et al., 2002; Klein et al., 2007; Goulson, 2010). There are more than 265 bumble bee species worldwide, 47 of which occur in the USA (Colla et al., 2011; Koch et al., 2012; Williams et al., 2014; Maebe et al., 2021). Despite their ecological and economic importance to wild and agricultural systems, bumble bee communities are undergoing drastic changes due to anthropogenic effects such as climate change and agricultural intensification and expansion (Kerr et al., 2015; Goulson et al., 2015; Fourcade et al., 2019; Kohler et al., 2020).

Global climate change has led to a rise in average temperatures, changes in precipitation patterns, and an increase in the frequency and intensity of extreme and localized weather events (Easterling et al., 2000; Meehl and Tebaldi, 2004). Changes in climate can have profound impacts on bumble bee species’ abundances, distributions, and population dynamics as well as overall community structure (Easterling et al., 2000; Parmesan, 2006; Kerr et al., 2015; Fourcade et al., 2019). Temperature is a pervasive selective pressure that influences bumble bee evolution and adaptation (Hines, 2008; Martinet et al., 2018; Williams et al., 2018; Pimsler et al. 2020). Given their annual colony cycle, bumble bees are exposed to a wide range of temperatures as they develop from spring to fall and gynes undergo winter diapause. Common throughout cool temperate, alpine, and arctic ecosystems, bumble bees can regulate their body temperature and generate heat in cold climates (Kerr et al., 2015; Pimsler et al., 2020), while also having adaptations to prevent overheating (i.e., wing fanning, thoracic or evaporative cooling) during the summer (Heinrich, 1976; Westhus et al., 2013). Despite these adaptations, frequent and intense heat waves can lead to increased mortality of bumble bees by inducing hyperthermic stress when foraging (for workers) or during nuptial behavior (for males) (Martinet et al., 2015; Oyen et al., 2016; Fourcade et al., 2019; Martinet et al., 2020). Additionally, precipitation has direct (e.g., desiccation) and indirect (e.g., floral resource availability) effects on bumble bee fitness, and can drive foraging periods and geographic ranges (Willmer and Stone, 1998; Williams et al., 2015; Jackson et al., 2018; Koch et al., 2019). Therefore, understanding how bioclimatic predictors (climate conditions related to species physiology) shape bumble bee assemblage composition (calculated as species richness and abundance) is of critical importance for evaluating climatic constraints. Climate change also interacts with other anthropogenic disturbances, such as land cover conversion and intensification, further altering species’ responses to these environmental conditions (Easterling et al., 2000; Kerr et al., 2015; Marshall et al., 2018; Fourcade et al., 2019).

Within agricultural systems, intensification and expansion of high-yielding field crops have led to extensive reductions in landscape composition (i.e., the number and amount of distinct land cover categories) and configuration (i.e., the spatial arrangement of those land cover categories), resulting in landscape simplification (Meehan et al., 2011; Pfeiffer et al., 2019; Nelson and Burchfield, 2021). Landscape simplification often reduces the availability, diversity, and distribution of nutritionally sufficient floral resources and nesting sites for bumble bees (Parys et al., 2021), leading them to be inadvertently extirpated in these agricultural environments (Westphal et al., 2003; Hall et al., 2017; Williams et al., 2012). While bumble bees can fly several kilometers to establish colonies and forage, they are also central place foragers with strong site fidelity (Goulson, 2010; Lepais et al., 2010; Rao and Strange, 2012; Ogilvie and Thomson, 2016). Therefore, increasing the diversity and connectedness of land use types that provide increased floral and nesting resources within areas surrounding agriculturally intensified lands can positively impact bumble bee assemblage composition and diversity (Kaiser-Bunbury et al., 2017; Miljanic et al., 2019).

While the individual effects of climate and various landscape factors have been studied, bumble bee research is just starting to evaluate the effects of multiple stressors simultaneously, underscoring the need to understand how bumble bees are affected by a range of co-occurring environmental changes across a range of geographic regions (Marshall et al., 2018; Fourcade et al., 2019; Escobedo-Kenefic et al., 2020; Kohler et al., 2020; Ganuza et al., 2022). Research incorporating land cover change with climate change continually reports that evaluating these factors together is important to better understand environmental change impacts on pollinators, which can then be used to implement effective conservation practices (Naeem et al., 2019; Kammerer et al., 2020). As bumble bees have strong species-specific associations with land use and climate variables, it is important to study both individual species responses and community responses to gain a more holistic understanding of how bumble bees will be impacted by environmental changes (Whitehorn et al., 2022). For example, imperiled *B. occidentalis* range-wide declines have been attributed to increasing temperatures during the warmest quarter of the year, severe drought years, and use of nitroguanidine neonicotinoid insecticides, while their occupancy is positively associated with increased forest and shrub area (Janousek et al., 2022). Meanwhile, previous research in Utah identified that bumble bee assemblage composition was highest at agricultural sites surrounded by more agricultural land cover, low temperatures, and high relative humidity, and lowest at agricultural sites with more urban land cover in the surrounding area, high temperatures, and low relative humidity (Christman et al., 2022a). Further, land use can amplify or offset changes in suitable habitat for bumble bees under various climate projections, emphasizing the importance of utilizing targeted land management practices on top of climate mitigation techniques to reduce the magnitude of bumble bee declines (Prestele et al., 2021). Overall, evaluating climate and land-use factors together is critical for managing pollinator populations and their environments.

In this study, we evaluated the effects of climate, landscape composition, and landscape configuration on bumble bee species assemblages in agricultural fields throughout nine states in the USA. Specifically, we tested bumble bee responses to climate and land use by modeling 1) occupancy (presence/absence); 2) species richness; 3) species-specific habitat requirements; and 4) assemblage-level responses and how those responses vary across geographic regions. Results from this study contribute to a better understanding of climate and landscape factors affecting bumble bees and their habitats throughout the USA. Detailed knowledge of species-specific relationships with climate and landscape variables across a geographic range is invaluable to improve targeted conservation and land management strategies to mitigate the effects of ongoing environmental changes.

## 2. MATERIALS AND METHODS

### 2.1. Overview

We combined richness, abundance, and distribution data for 18 bumble bee species with bioclimatic, landscape composition, and landscape configuration data to evaluate the effects of multiple environmental stressors on bumble bee occupancy and species assemblages throughout the USA. Bumble bees were collected as bycatch within pest monitoring traps placed in 433 agricultural fields throughout nine states from 2018 to 2020. Bioclimatic data was derived at a 1 km spatial scale using historical Daymet monthly precipitation, minimum temperature, and maximum temperature data from 2000 to 2020 (Thornton et al., 2020). Landscape composition and configuration data was derived at a 1 km spatial scale from land cover values obtained from the USDA National Agricultural Statistics Service CropScape and Cropland Data Layer from 2018 to 2020 (USDA NASS CDL, 2018–2020). A logistic regression, a generalized additive mixed model, a canonical correspondence analysis, and four multivariate regression trees were conducted to comprehensively assess bumble bee occupancy and assemblage-level responses in relation to climate and land use variables. All data were analyzed using R version 4.3.1 (R Core Team, 2022). Details for all methodological and analytical steps are described in detail in the following sections.

### 2.2. Collection of Bumble Bees

Bumble bees were collected within pest monitoring traps that were placed by state cooperators in agricultural fields across diverse regions in the USA as part of early-detection surveys for invasive lepidopterans following Spears et al., (2016) and U.S. Department of Agriculture, Animal and Plant Health Inspection Service, Cooperative Agricultural Pest Survey approved methods for pest surveillance (CAPS, 2022). Previous research identified that bumble bees are attracted to pest monitoring traps and suggested that these captures be used to advance knowledge of biodiversity, population fluctuations, and other ecological objectives (Buchholz et al., 2011; Spears and Ramirez, 2015; Sipolski et al., 2019; Whitfield et al., 2019; Grocock et al., 2020; Grocock and Evenden, 2020; Parys et al., 2021; Spears et al., 2016, 2021; Christman et al., 2022a).

This study included a total of 433 agricultural fields throughout Florida, Indiana, Kansas, Kentucky, Maryland, South Carolina, Utah, Virginia, and West Virginia from 2018 to 2020, where the number of sites varied by state, year, and target pest (Table 1). Target pests included Christmas berry webworm (CBW, *Cryptoblabes gnidiella* Milliere, 1867), cotton cutworm (CC, *Spodoptera litura* Fabricius, 1775), Egyptian cottonworm (EC, *Spodoptera littoralis* Boisduval, 1833), golden twin spot moth (GTS, *Chrysodeixis chalcites* Esper, 1789), Old World bollworm (OWB, *Helicoverpa armigera* Hübner, 1808), and silver Y moth (SYM, *Autographa gamma* Linnaeus, 1758). Following methodology outlined in Christman et al. (2022a) and CAPS (2022), multi-colored (green canopy, yellow funnel, and white bucket) bucket traps (International Pheromone Systems, Cheshire, UK) were placed 20 m apart and hung 1.5 m above the ground along the edge of vegetable or other commodity crop fields (e.g., alfalfa, corn, small grain) as part of the CAPS monitoring program. Each trap contained a pheromone lure for a single target pest inside the lure basket of the trap canopy. An insecticide strip (Hercon Vaportape II: 10% dimethyl 2,2-dichlorovinyl phosphate, Hercon Environmental Corporation, Emigsville, PA) and a small, cellulose sponge were placed inside each bucket to kill the captured insects and absorb rainwater, respectively. Insecticide strips and pheromone lures for CBW, GTS, OWB, and SYM were replaced no more than every 28 days, whereas pheromone lures for CC and EC were changed no more than every 84 days. Although the collection period for traps varied by state, most traps were serviced biweekly (monthly in Kentucky) from May to August, but some states had an extended trapping season based on the period of expected pest activity (Table 1). Since lure comparisons were not the intent of this study (but see Spears et al., 2016 for lure-specific analyses), trap data were combined by study site and collection period.

**Table 1.** The number of sites, target pests, and collection period by state and year. Target pests included Christmas berry webworm (CBW), cotton cutworm (CC), Egyptian cottonworm (EC), golden twin spot moth (GTS), Old World bollworm (OWB), and silver Y moth (SYM).

| State | Number of Sites | Target Pest(s) | Collection Period |
| --- | --- | --- | --- |
| *2018* |  |  |  |
| Kansas | 86 | CC, EC | July-October |
| Utah | 30 | CC, EC, OWB | April-September |
| West Virginia | 9 | CC, EC, GTS, OWB, SYM | June-September |
| *2019* |  |  |  |
| Florida | 16 | OWB | April-September |
| Indiana | 6 | CC, EC, GTS, OWB, SYM | May-August |
| Kentucky | 41 | CC, EC, GTS, OWB, SYM | May-October |
| Maryland | 26 | CC, OWB, SYM | June-July |
| South Carolina | 16 | CC, EC, OWB | June-October |
| Utah | 30 | CC, EC, OWB | June-August |
| Virginia | 13 | OWB | July-September |
| West Virginia | 32 | CBW, CC, EC, GTS, OWB, SYM | May-October |
| *2020* |  |  |  |
| Indiana | 6 | CC, EC, GTS, OWB, SYM | April-August |
| Kentucky | 63 | CC, EC, GTS, OWB, SYM | May-September |
| Utah | 30 | CC, EC, OWB | May-August |
| Virginia | 12 | OWB | August-September |
| West Virginia | 17 | CBW, CC, EC, OWB, SYM | May-September |

Trap contents were screened for target pests by state cooperators, and all non-target captures (bycatch) were sent to the Utah State University Department of Biology. Bumble bees were separated from all other non-target specimens and then stored in a freezer at -18°C until they could be pin-mounted, labeled, and identified to species using taxonomic keys (Colla et al., 2011; Koch et al., 2012; Williams et al., 2014). All specimens were deposited into a collection in the Department of Biology at Utah State University in Logan, Utah, USA, and records were entered into the USDA, Agricultural Research Service, Pollinating Insect – Biology, Management, and Systematics Research Unit, National Pollinating Insects Database.

### 2.3. Bioclimatic Variables

Historical weather data over the past 20 years (2000–2020) were extracted from each site at a 1 km spatial resolution for averaged monthly precipitation, minimum temperature, and maximum temperature using Daymet monthly climate summaries (Thornton et al., 2020). Nineteen bioclimatic variables from WorldClim were derived from the monthly precipitation and temperature values to generate more biologically meaningful variables with the *dismo* library (Fick and Hijmans, 2017; Hijmans et al., 2020). Using an expert based selection process, this subset was reduced further by selecting six variables that represent annual climatic trends and extremes: annual mean temperature, maximum temperature of warmest month, minimum temperature of coldest month, annual precipitation, precipitation of wettest month, and precipitation of driest month (Fick and Hijmans, 2017; Hijmans et al., 2020; Jackson et al., 2020).

### 2.4. Landscape Composition and Configuration

The elevation of each site was extracted from the North American Elevation 1 km Resolution GRID (U.S. Department of the Interior, 2021). Land cover values from 2018 to 2020 were obtained from USDA National Agricultural Statistics Service (NASS) CropScape and Cropland Data Layer (CDL) (USDA NASS CDL, 2018–2020), which maps data at a 30-meter spatial resolution.

The 255 Cropland Data Layer land cover classes were aggregated into five land cover categories: bumble bee attractive crops, bumble bee unattractive crops, rangeland, forest, and developed land (Supp. Table 1). The attractiveness of agricultural crops to bumble bees was determined using the Pollinator Attractiveness Crop List produced by the USDA (2017). Bumble bee attractive crops included plants such as tomatoes, peppers, berries, and alfalfa, while bumble bee unattractive crops included plants such as corn, sorghum, and wheat. Here, we note that in this study, alfalfa was included as a bumble bee attractive crop since it can flower up to 25% prior to being harvested, providing a pulse of resources for bumble bees (USDA, 2017). Rangeland referred to lands suitable for grazing or browsing, including switchgrass, shrubland, wetlands (woody and herbaceous), grasslands, and pastures. Forest included deciduous, evergreen, and mixed forests. Developed land referred to urban environments that have been built up with non-impervious surface cover, including low, medium, and high intensity lands. The number of pixels of each land cover category was then extracted from a 1 km buffer surrounding each site and the percentage of bumble bee attractive crops, bumble bee unattractive crops, rangeland, forest, and developed land was quantified to determine the landscape composition surrounding our surveyed agricultural sites.

Additionally, landscape configuration indices: contiguity, and interspersion and juxtaposition, were calculated at a 1 km buffer surrounding each site using the 255 Cropland Data Layer land cover classes with the *landscapemetric* library (Hesselbarth et al., 2019). Contiguity refers to the spatial connectedness of land cover classes within a landscape. Interspersion and juxtaposition refer to the arrangement, relationship, and proximity of different land cover classes in a landscape (Hesselbarth et al., 2019).

### 2.5. Data Analysis

Four aspects of bumble bee species composition were measured for each state: total count, richness (number of species), Pielou’s evenness (abundance per species), and Shannon diversity (which accounts for evenness and richness) with the *vegan* and *codyn* libraries. The weekly bumble bee collection rate for each state was quantified each year to standardize differences between state collection periods. Bubble maps were used to visualize bumble bee distribution, abundance, and diversity throughout the surveyed states.

Variance inflation factor was used to test for multicollinearity between the selected bioclimatic and landscape variables. Variables with a variance inflation factor greater than 10 were removed in descending order until all values were lower than 10 to reduce collinearity between the explanatory variables. Maximum temperature of warmest month, minimum temperature of coldest month, precipitation of wettest month, precipitation of driest month, elevation, bumble bee attractive crops, bumble bee unattractive crops, forest, developed land, contiguity, and interspersion and juxtaposition were included as the bioclimatic variables and landscape indices within the following models.

A logistic regression was used to identify which landscape and bioclimatic variables were predictive of bumble bee occupancy (presence/absence) in a landscape. Stepwise model selection by AIC was performed on the full logistic regression model to select the best set of predictor variables. A final logistic regression was then conducted with the resulting best set of selected landscape and bioclimatic variables that accounted for bumble bee presences and absences. Odds ratios were calculated to indicate the strength and direction of association between bumble bee presence and absence and each of the individual landscape and bioclimatic variables.

A generalized additive mixed model was used to describe bumble bee species richness in relation to linear and non-linear bioclimatic and landscape variables among all surveyed sites while incorporating spatial and temporal autocorrelation residuals with the *mcgv* and *nlme* libraries. All predictor variables were initially smoothed using p-splines to account for non-linearities. Variables with an effective degree of freedom of 1 suggested the term was reduced to a simple linear effect, and thus did not need to be smoothed.

A canonical correspondence analysis (CCA) was used identify species-specific habitat requirements by assessing correlations among explanatory variables (bioclimatic variables and landscape indices) and response variables (*Bombus* species assemblages) from 2018 to 2020 with the *vegan* and *picante* libraries. Variance inflation factor was used again to test for multicollinearity in the CCA. Precipitation of driest month was removed to reduce collinearity between the explanatory variables. A permutation test was used to determine the significance of each variable and the overall model.

Multivariate regression trees were used to document assemblage-level responses by evaluating the interactions between bumble bee species abundance and environmental variables by geographic region from 2018 to 2020 with the *mvpart* library. Study sites were split into one of four USDA Farm Production Regions: corn belt/Appalachian/northeast, southeast, northern plains, and mountains. Corn belt/Appalachian/northeast included sites within Indiana, Kentucky, Maryland, Virginia, and West Virginia (n = 225). Southeast included sites within Florida and South Carolina (n = 32). Northern plains included sites within Kansas (n = 86), and mountains included sites within Utah (n = 90). The multivariate regression tree groups sites based on repeated splits in environmental variable values, minimizing dissimilarity within site groups. Each leaf represents species assemblages and the environmental variable values associated with the sites, which are displayed in the form of a tree. A cross-validation with 100 iterations was generated to identify the overall best-fit predictive tree (± 1 standard error) and to evaluate the predictive ability of each multivariate regression tree.

## 3. RESULTS

### 3.1. Collection of Bumble Bees

From 2018 to 2020, a total of 5,004 bumble bees representing 18 species were collected across nine states (Table 2). Collection rates varied by state and year. For example, Florida had extremely low collection rates of one bumble bee per week in 2019, whereas over forty bumble bees were collected per week in Utah (2018 and 2020) and West Virginia (2019) (Table 3). *Bombus fervidus* (Fabricus, 1798)*, B. bimaculatus* (Cresson, 1863), *B. impatiens* (Cresson, 1863), *B. pensylvanicus* (De Geer, 1773), and *B. huntii* (Greene, 1860) were the five most abundant species within traps, comprising 84% of total captures (Fig. 1). Bumble bee species diversity was consistently highest in Indiana, Kentucky, Utah, and West Virginia (Fig. 2, Table 3).

**Figure 1.**
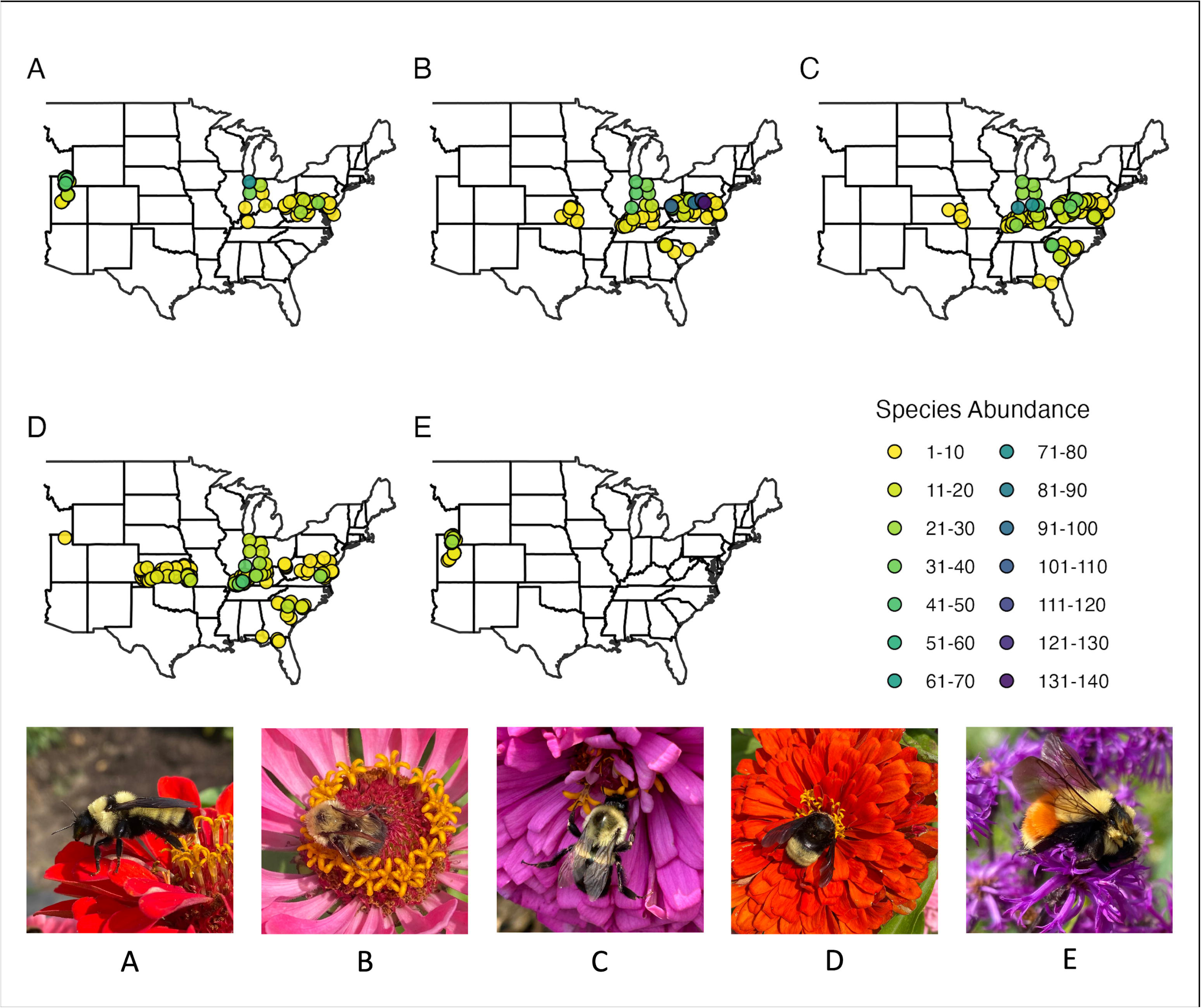
Bubble map showing the distribution and abundance of the five most abundant *Bombus* species: (A) *B. fervidus,* (B) *B. bimaculatus,* (C) *B. impatiens,* (D) *B. pensylvanicus,* and (E) *B. huntii* throughout nine states in the USA from 2018 to 2020. Different colors correspond to different levels of species abundance.

**Figure 2.**
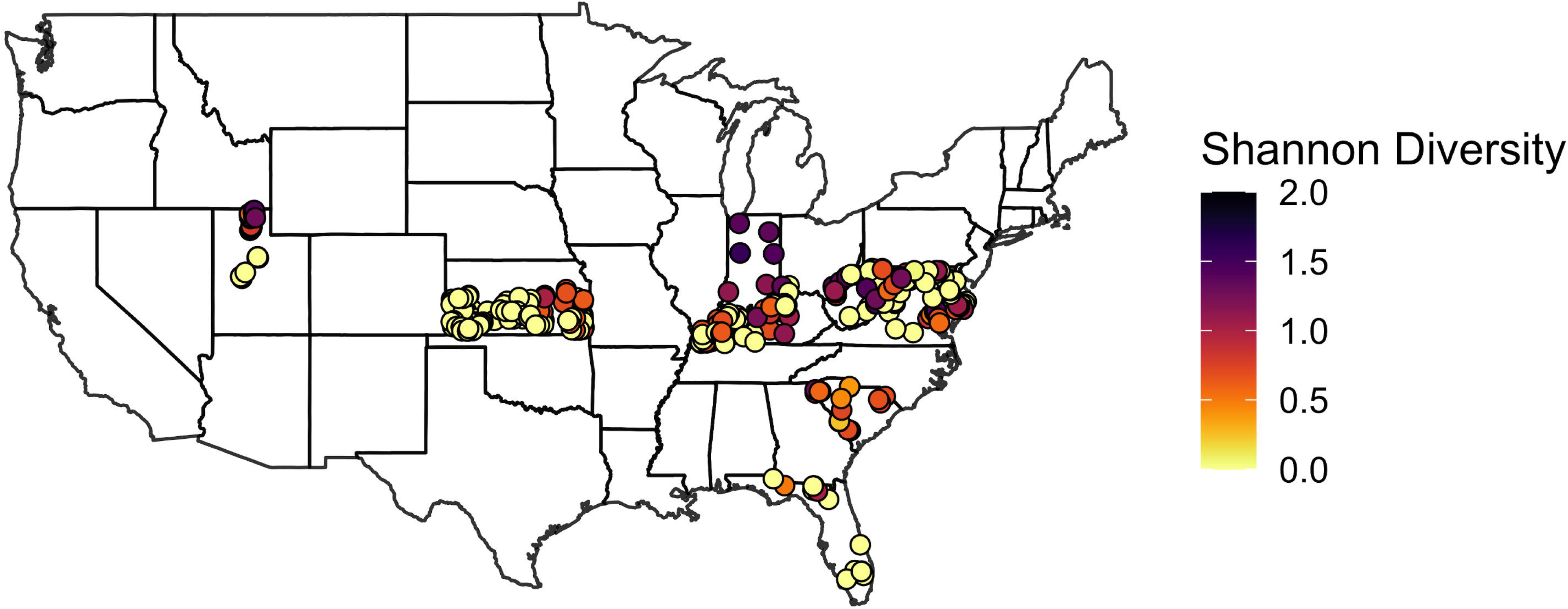
Bubble map visualizing bumble bee Shannon diversity throughout nine states in the USA from 2018 to 2020. Different colors correspond to different levels of Shannon diversity.

**Table 2.**
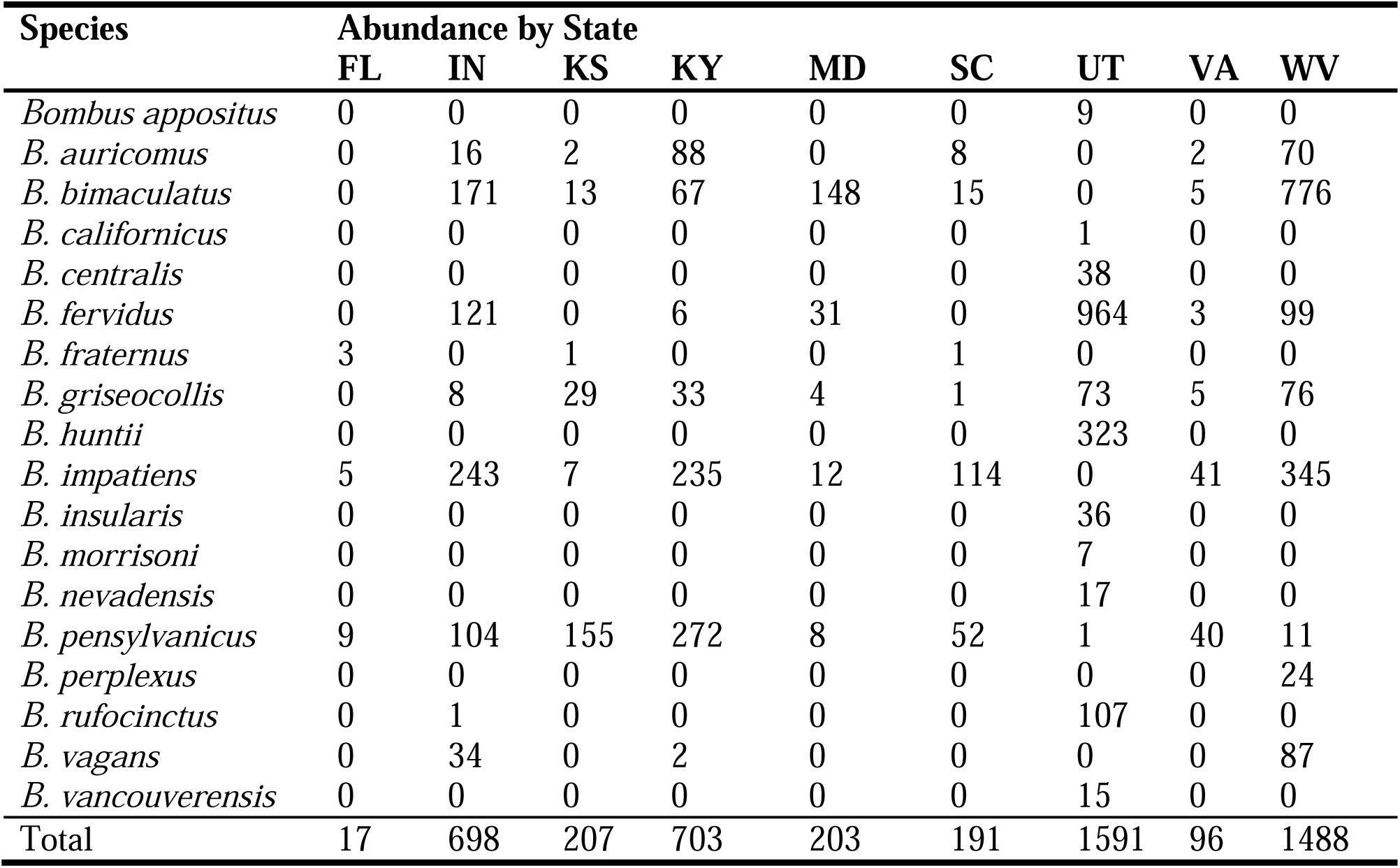
Bumble bee assemblage composition (species richness and abundance) in Florida, Indiana, Kansas, Kentucky, Maryland, South Carolina, Utah, Virginia, and West Virginia from 2018 to 2020.

| Species | Abundance by State |  |  |  |  |  |  |  |  |
| --- | --- | --- | --- | --- | --- | --- | --- | --- | --- |
|  | FL | IN | KS | KY | MD | SC | UT | VA | WV |
| *Bombus appositus* | 0 | 0 | 0 | 0 | 0 | 0 | 9 | 0 | 0 |
| *B. auricomus* | 0 | 16 | 2 | 88 | 0 | 8 | 0 | 2 | 70 |
| *B. bimaculatus* | 0 | 171 | 13 | 67 | 148 | 15 | 0 | 5 | 776 |
| *B. californicus* | 0 | 0 | 0 | 0 | 0 | 0 | 1 | 0 | 0 |
| *B. centralis* | 0 | 0 | 0 | 0 | 0 | 0 | 38 | 0 | 0 |
| *B. fervidus* | 0 | 121 | 0 | 6 | 31 | 0 | 964 | 3 | 99 |
| *B. fraternus* | 3 | 0 | 1 | 0 | 0 | 1 | 0 | 0 | 0 |
| *B. griseocollis* | 0 | 8 | 29 | 33 | 4 | 1 | 73 | 5 | 76 |
| *B. huntii* | 0 | 0 | 0 | 0 | 0 | 0 | 323 | 0 | 0 |
| *B. impatiens* | 5 | 243 | 7 | 235 | 12 | 114 | 0 | 41 | 345 |
| *B. insularis* | 0 | 0 | 0 | 0 | 0 | 0 | 36 | 0 | 0 |
| *B. morrisoni* | 0 | 0 | 0 | 0 | 0 | 0 | 7 | 0 | 0 |
| *B. nevadensis* | 0 | 0 | 0 | 0 | 0 | 0 | 17 | 0 | 0 |
| *B. pensylvanicus* | 9 | 104 | 155 | 272 | 8 | 52 | 1 | 40 | 11 |
| *B. perplexus* | 0 | 0 | 0 | 0 | 0 | 0 | 0 | 0 | 24 |
| *B. rufocinctus* | 0 | 1 | 0 | 0 | 0 | 0 | 107 | 0 | 0 |
| *B. vagans* | 0 | 34 | 0 | 2 | 0 | 0 | 0 | 0 | 87 |
| *B. vancouverensis* | 0 | 0 | 0 | 0 | 0 | 0 | 15 | 0 | 0 |
| Total | 17 | 698 | 207 | 703 | 203 | 191 | 1591 | 96 | 1488 |

**Table 3.** Community composition indices of bumble bees collected in Florida, Indiana, Kansas, Kentucky, Maryland, South Carolina, Utah, Virginia, and West Virginia from 2018 to 2020.

| **State** | **Total Count** | **Richness** | **Pielou's Evenness** | **Shannon Diversity** | **Collection Rate by Week** |
| --- | --- | --- | --- | --- | --- |
| *2018* |  |  |  |  |  |
| Kansas | 207 | 6 | 0.47 | 0.85 | 14.21 |
| Utah | 746 | 11 | 0.49 | 1.18 | 47.47 |
| West Virginia | 237 | 8 | 0.69 | 1.43 | 19.07 |
| *2019* |  |  |  |  |  |
| Florida | 17 | 3 | 0.91 | 1.00 | 1.37 |
| Indiana | 310 | 7 | 0.79 | 1.54 | 25.53 |
| Kentucky | 535 | 7 | 0.71 | 1.37 | 32.01 |
| Maryland | 40 | 5 | 0.92 | 1.47 | 11.20 |
| South Carolina | 191 | 6 | 0.59 | 1.05 | 12.38 |
| Utah | 217 | 9 | 0.45 | 0.98 | 22.34 |
| Virginia | 103 | 6 | 0.65 | 1.17 | 14.71 |
| West Virginia | 932 | 8 | 0.71 | 1.47 | 48.33 |
| *2020* |  |  |  |  |  |
| Indiana | 388 | 8 | 0.78 | 1.62 | 23.82 |
| Kentucky | 168 | 6 | 0.78 | 1.40 | 7.30 |
| Utah | 594 | 11 | 0.56 | 1.34 | 51.98 |
| Virginia | 10 | 3 | 0.58 | 0.64 | 2.00 |
| West Virginia | 309 | 8 | 0.57 | 1.18 | 18.18 |

### 3.2. Bumble Bee Presence-Absence Characterized by Climate and Landscape

Maximum temperature of warmest month, precipitation of wettest month, elevation, bumble bee attractive crops, bumble bee unattractive crops, contiguity, and interspersion and juxtaposition were selected as the best set of variables in the logistic regression that accounted for bumble bee occupancy (presences/absences) in a landscape (Table 4). Bumble bee presence within a landscape was positively associated with bumble bee attractive crops. For every 1% increase in bumble bee attractive crops, there was a corresponding 1.02x increase in the odds that bumble bees were present (Table 4). Meanwhile, maximum temperature of warmest month and bumble bee unattractive crops were associated with the absence of bumble bees in a landscape. For every 1°C increase in maximum temperature of warmest month, there was a corresponding 0.85x decrease in the odds the bumble bees were present. For every 1% increase in bumble bee unattractive crops, the odds that bumble bees were present decreased by 0.99x (Table 4). Precipitation of wettest month, elevation, contiguity, and interspersion and juxtaposition were not significant, but were retained for overall model fit.

**Table 4.** Logistic regression model results for the best set of selected bioclimatic and landscape variables that accounted for bumble bee presences and absences based on stepwise model

| **Predictor Variable** | **Odds Ratio** | **95% CI** | **Z-Score** | ***p*-value** |
| --- | --- | --- | --- | --- |
| Maximum Temperature of Warmest Month | 0.852 | (0.736, 0.979) | -2.210 | 0.027 |
| Precipitation of Wettest Month | 0.958 | (0.907, 1.012) | -1.531 | 0.126 |
| Elevation | 1.001 | (1.000, 1.001) | 1.958 | 0.050 |
| Bumble Bee Attractive Crops | 1.019 | (1.002, 1.037) | 2.201 | 0.028 |
| Bumble Bee Unattractive Crops | 0.987 | (0.975, 0.998) | -2.252 | 0.024 |
| Contiguity | 1.053 | (0.993, 1.119) | 1.680 | 0.093 |
| Interspersion and Juxtaposition | 1.032 | (0.999, 1.068) | 1.858 | 0.063 |

### 3.3. Climate and Landscape Impact on Bumble Bee Species Richness

Bumble bee species richness was significantly associated with maximum temperature of warmest month, minimum temperature of coldest month, precipitation of wettest month, elevation, bumble bee attractive crops, and contiguity, while accounting for spatial and temporal covariance among the surveyed sites in the generalized additive mixed model (Fig. 3, Table 5, and Table 6). Based on partial effects plots, bumble bee species richness increased as maximum temperature of warmest month approached 29°C and then declined with variation as temperatures rose to 36°C (Fig. 3A). Meanwhile, species richness declined as minimum temperature of coldest month approached -5°C and then increased slightly as temperatures approached 10°C (Fig. 3B). As precipitation of wettest month increased, bumble bee species richness decreased (Fig. 3C). Bumble bee species richness steadily increased with higher elevations and as bumble bee attractive crops became more prominent in the landscape (Fig. 3D & 3E). Bumble bee species richness increased slightly as contiguity approached 0.2 and then declined as it approached 0.3 (Fig. 3F).

**Figure 3.**
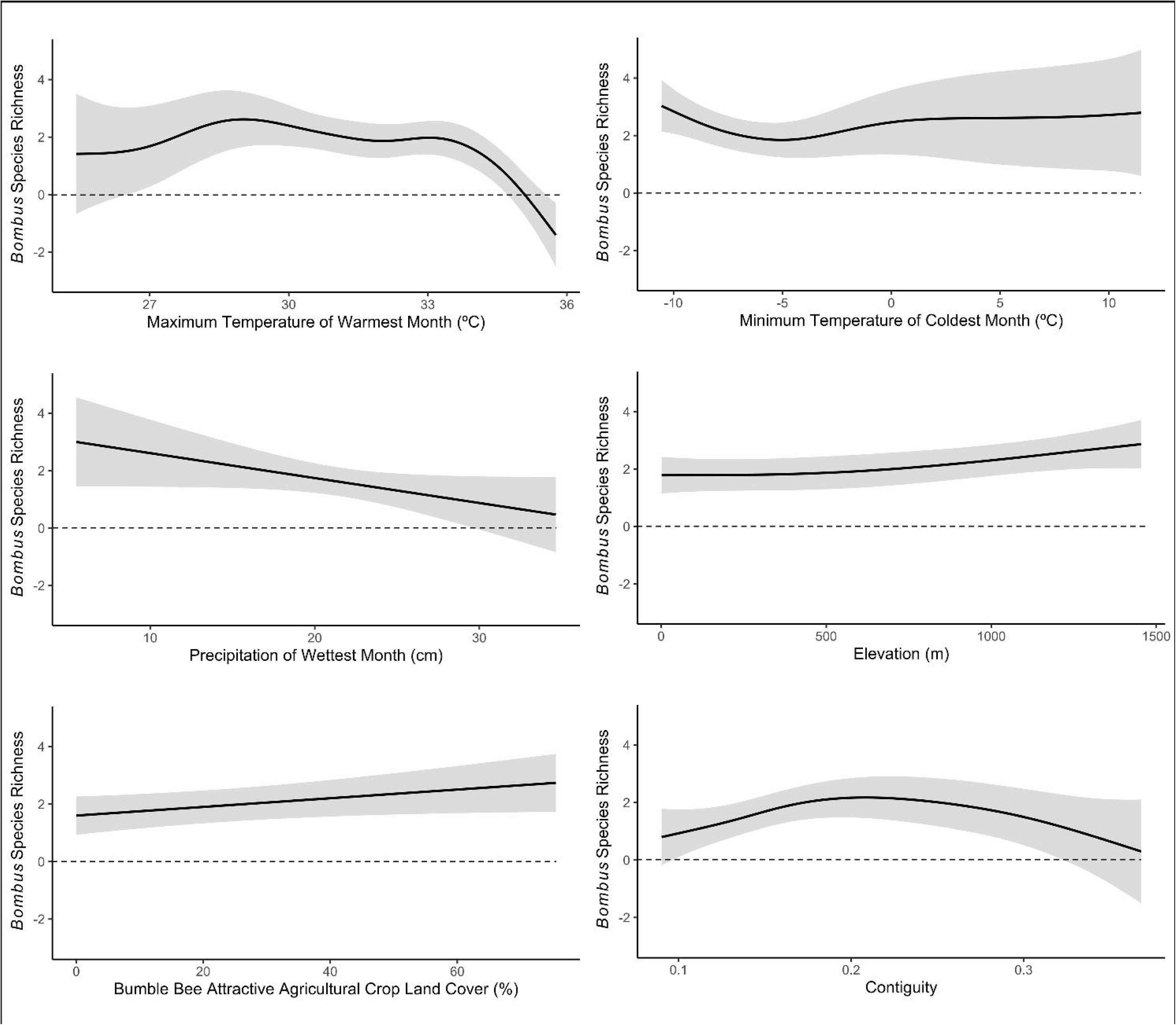
Generalized additive model plots showing the partial effects of A) maximum temperature of warmest month, B) minimum temperature of coldest month, C) precipitation of wettest month, D) elevation, E) bumble bee attractive crops, and F) contiguity on bumble bee species richness. Gray areas represent confidence intervals ± 2 standard errors.

**Table 5.** Generalized additive mixed model results describing bumble bee species richness in relation to the non-linear bioclimatic and landscape variables, while accounting for spatial and temporal autocorrelation. Model results include the effective degrees of freedom, *F-*values, and *p-*values for each of the p-spline smoothed (non-linear) effects.

| **Smoothed Predictor Variable** | **EDF** | ***F*-value** | ***p*-value** |
| --- | --- | --- | --- |
| Maximum Temperature of Warmest Month | 5.178 | 8.994 | 0.001 |
| Minimum Temperature of Coldest Month | 2.891 | 3.873 | 0.037 |
| Contiguity | 2.599 | 4.277 | 0.011 |

**Table 6.** Generalized additive mixed model results describing bumble bee species richness in relation to the linear bioclimatic and landscape variables, while accounting for spatial and temporal autocorrelation. Model results include the coefficient estimate, standard error, and *p*-

| **Linear Predictor Variable** | **Coefficient Estimate** | **Std. Error** | ***p*-value** |
| --- | --- | --- | --- |
| Precipitation of Wettest Month | -0.107 | 0.043 | 0.014 |
| Precipitation of Driest Month | 0.273 | 0.175 | 0.119 |
| Elevation | 0.001 | 0.000 | 0.049 |
| Bumble Bee Attractive Crops | 0.017 | 0.008 | 0.034 |
| Bumble Bee Unattractive Crops | -0.012 | 0.007 | 0.080 |
| Forest | 0.002 | 0.007 | 0.796 |
| Developed Land | -0.008 | 0.009 | 0.375 |
| Interspersion and Juxtaposition | 0.004 | 0.013 | 0.748 |

### 3.4. Bumble Bee Habitat Associations

The permutation test determined that the overall canonical correspondence analysis model was statistically significant (F_10, 293_ = 18.31, *p*-value = 0.001). Additionally, the permutation test by term (i.e., explanatory variables) determined that bumble bee abundances were significantly correlated with maximum temperature of warmest month, minimum temperature of coldest month, precipitation of wettest month, elevation, bumble bee attractive crops, bumble bee unattractive crops, forest, developed land, and interspersion and juxtaposition (Table 7). Over the three-year study period, these variables explained 35.99% of variation in bumble bee assemblages. *Bombus appositus* (Cresson, 1878)*, B. californicus* (Smith, 1854)*, B. centralis* (Cresson, 1864)*, B. fervidus* (Fabricius, 1798)*, B. huntii* (Greene, 1860)*, B. insularis* (Smith, 1861)*, B. morrisoni* (Cresson, 1878)*, B. nevadensis* (Cresson, 1874)*, B. rufocinctus* (Cresson, 1863), and *B. vancouverensis* (Cresson, 1878) were clustered together (Fig. 4). These western species shared similar habitat requirements, particularly high elevation environments with high proportions of bumble bee attractive and unattractive crops within the surrounding area, high maximum temperatures of warmest month, and increased interspersion and juxtaposition. Meanwhile, all eastern bumble bee species were associated with low elevation environments (Fig. 4). Additionally, *B. auricomus* (Robertson, 1903)*, B. fraternus* (Smith, 1854)*, B. impatiens* (Cresson, 1863), and *B. pensylvanicus* (De Geer, 1773) De Geer, 1773) were associated with high minimum temperatures of coldest month and increased precipitation of wettest month (Fig. 4). Further, *B. bimaculatus* (Cresson, 1863)*, B. perplexus* (Cresson, 1863), and *B. vagans* (Smith, 1854) were associated with habitats with high values of forest in the surrounding area. *Bombus griseocollis* (De Geer, 1773) was not associated with high or low values of any of the environmental variables, meaning they were found ubiquitously throughout the habitats regardless of the bioclimatic variable, landscape composition and configuration indice values (Fig. 4). Given the distinct groupings observed in the canonical correspondence analysis, we then evaluated bumble bee habitat associations by geographic region.

**Figure 4.**
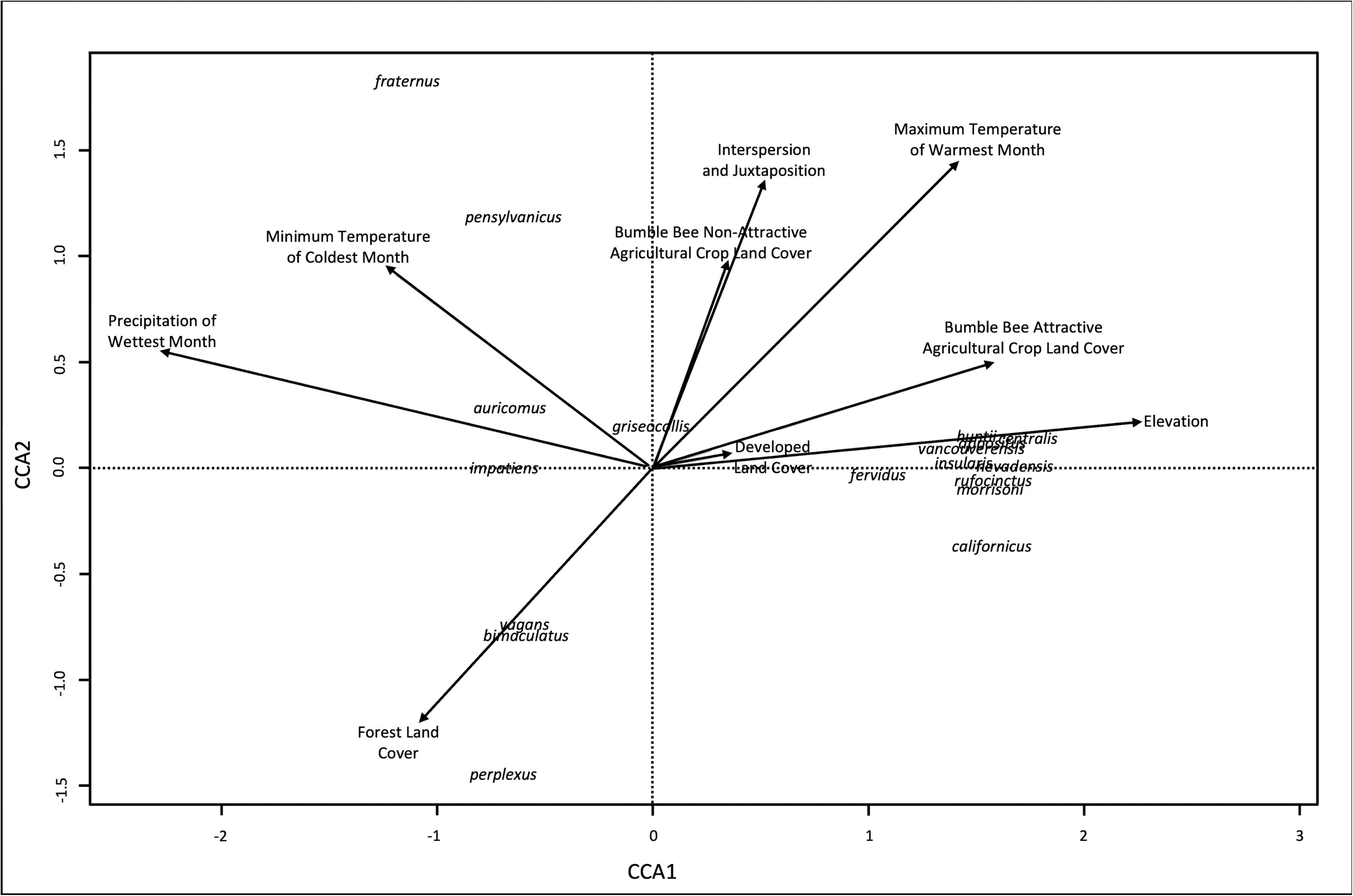
Canonical correspondence analyses of the *Bombus* assemblage data in relation to environmental variables (indicated by arrows) from 2018 to 2020.

**Table 7.**
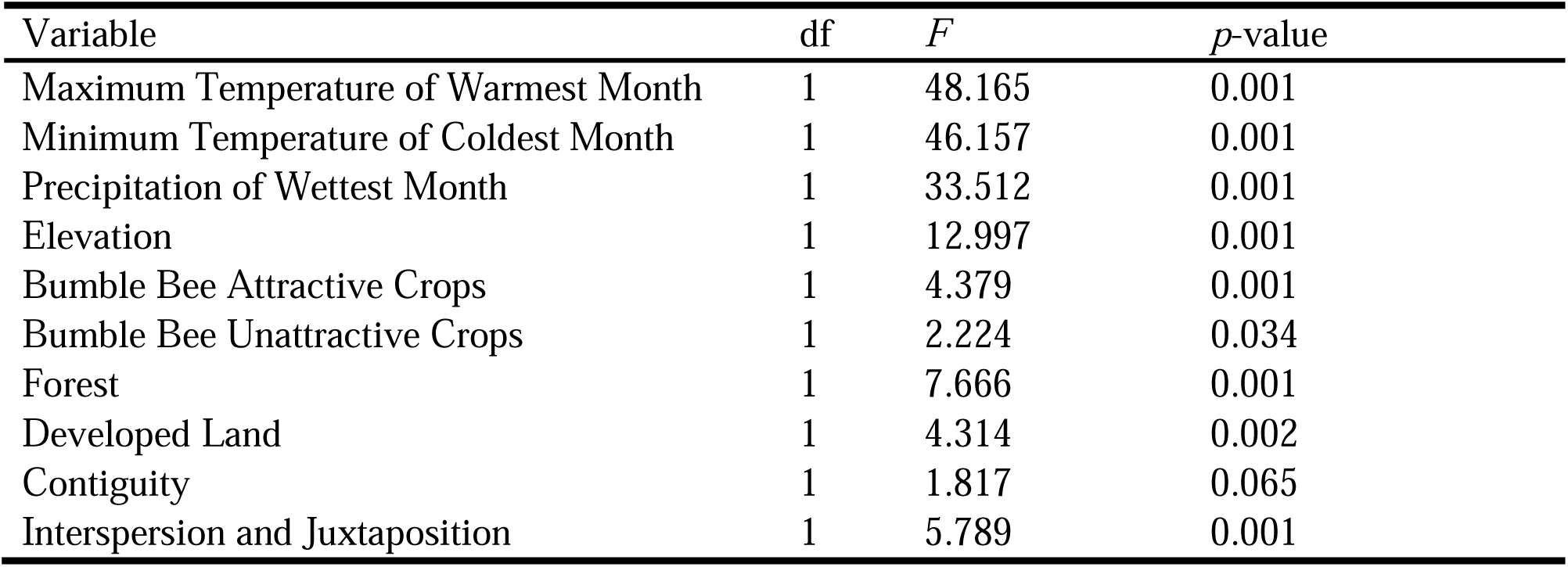
Significance of each explanatory variable from 2018 to 2020 based on a permutation test for the canonical correspondence analysis model.

| Variable | df | *F* | *p*-value |
| --- | --- | --- | --- |
| Maximum Temperature of Warmest Month | 1 | 48.165 | 0.001 |
| Minimum Temperature of Coldest Month | 1 | 46.157 | 0.001 |
| Precipitation of Wettest Month | 1 | 33.512 | 0.001 |
| Elevation | 1 | 12.997 | 0.001 |
| Bumble Bee Attractive Crops | 1 | 4.379 | 0.001 |
| Bumble Bee Unattractive Crops | 1 | 2.224 | 0.034 |
| Forest | 1 | 7.666 | 0.001 |
| Developed Land | 1 | 4.314 | 0.002 |
| Contiguity | 1 | 1.817 | 0.065 |
| Interspersion and Juxtaposition | 1 | 5.789 | 0.001 |

### 3.5. Climate and Landscape Impact on Bumble Bee Assemblages by Geographic Region

The multivariate regression tree for the corn belt/Appalachian/northeast geographic region resulted in a five-leaf tree where branching was determined by developed land, minimum temperature of coldest month, and maximum temperature of warmest month (Error = 0.484, CV Error = 0.979, SE = 0.473) (Fig 5). *Bombus bimaculatus* was the predominant species collected across all leaves. Sites with higher developed land and higher minimum temperature of coldest month had the most individual bumble bees present, with 1,153 bumble bees across seven species (Fig. 5A). Meanwhile, sites with higher developed land, moderate minimum temperature of coldest month, and higher maximum temperature of warmest month had the highest species richness, with 880 bumble bees collected across nine species (Fig. 5B). Sites with higher developed land, lower minimum temperature of coldest month, and higher maximum temperature of warmest month had lower abundances of the seven bumble bee species collected (Fig. 5C). Further, 761 bumble bees from 8 species were found at sites with higher developed land, lower minimum temperature of coldest month, and lower maximum temperature of warmest month (Fig. 5D). Bumble bee assemblages were the least rich in terms of richness and abundance at sites with lower proportions of developed land (Fig. 5E).

**Figure 5.**
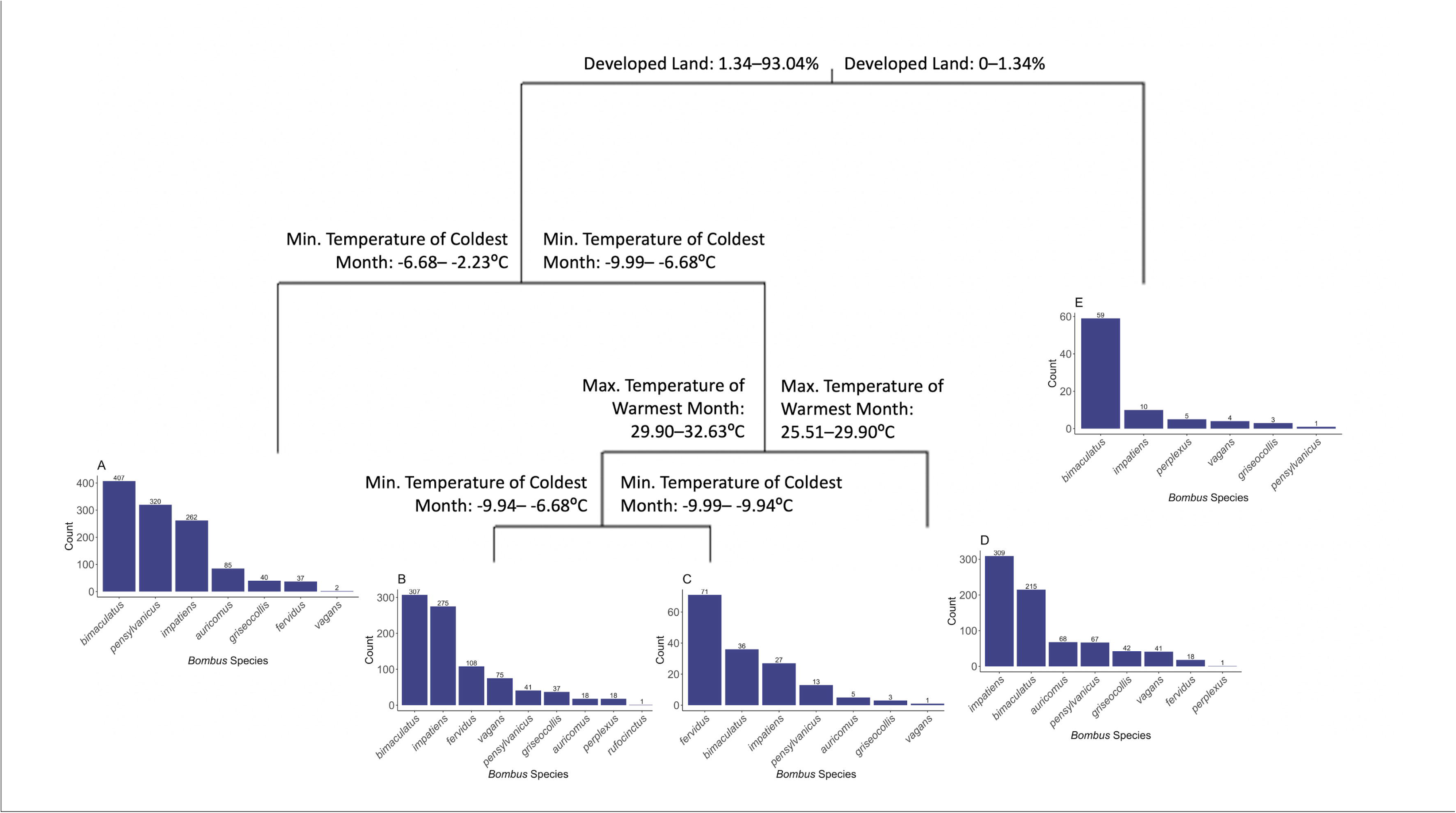
Multivariate regression tree for the corn belt/Appalachian geographic region, which resulted in branching at developed land, minimum temperature of coldest month, and maximum temperature of warmest month. The five leaves (indicated with letters under each branch) represent bumble bee species assemblages (richness and abundance) and the environmental variable values associated with the study sites.

The multivariate regression tree for the southeast geographic region resulted in a four-leaf tree where branching was determined by forest land, precipitation of wettest month, and maximum temperature of warmest month (Error = 0.278, CV Error = 1.63, SE = 0.572) (Fig 6). No bumble bees were found at sites with lower forest land, higher precipitation of wettest month, and lower maximum temperature of warmest month (Fig. 6A). Only one *B. pensylvanicus* was found at sites with lower forest land, higher precipitation of wettest month, and higher maximum temperature of warmest month (Fig. 6B). Meanwhile, sites with lower forest land and lower precipitation of wettest month had 97 bumble bees across four species (Fig. 6C). Bumble bee assemblages were most rich at sites with higher forest land, where 110 bumble bees across six species were found at these sites (Fig. 6D).

**Figure 6.**
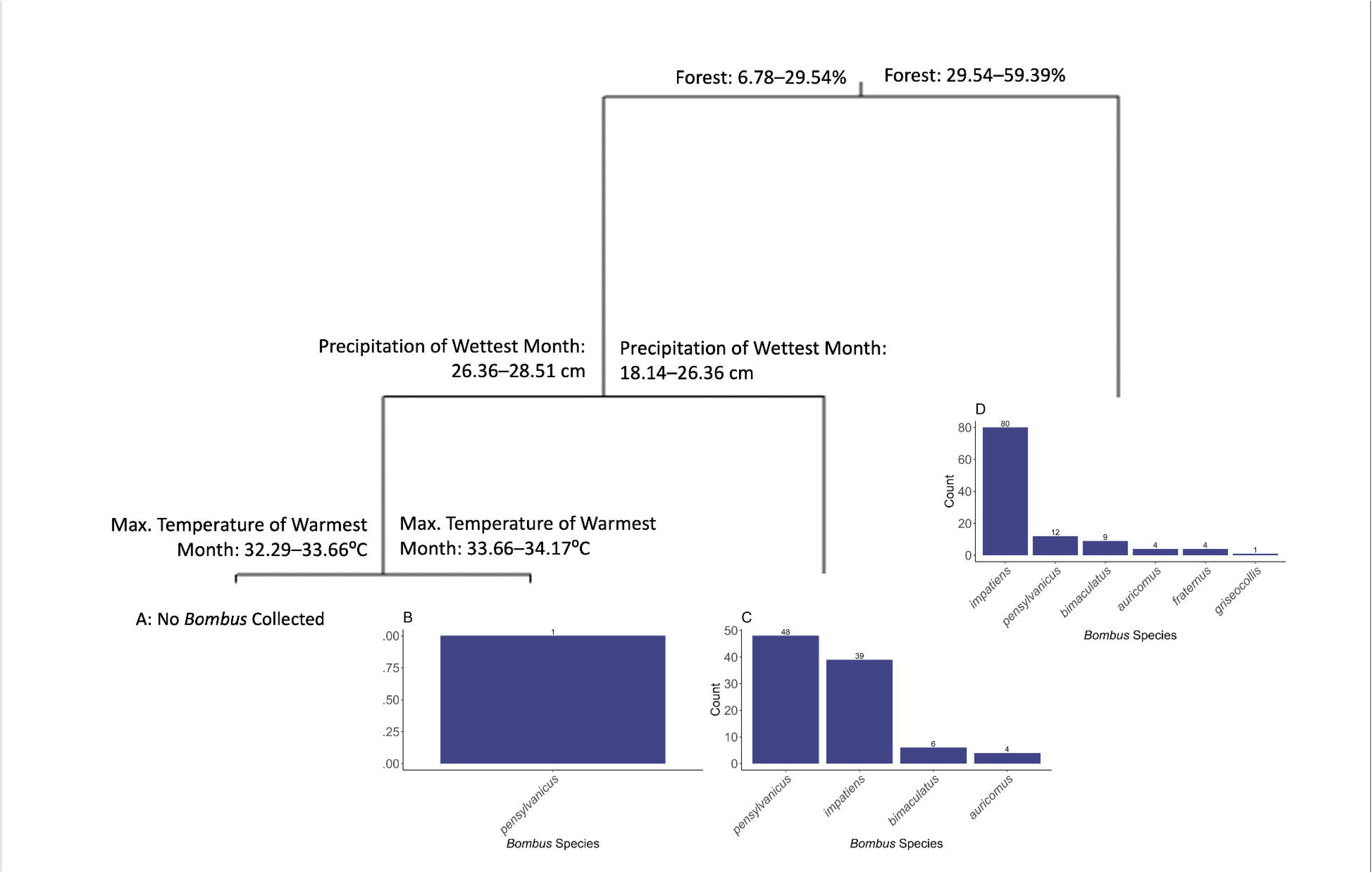
Multivariate regression tree for the southeast geographic region, which resulted in branching at forests, precipitation of wettest month, and maximum temperature of warmest month. The four leaves (indicated with letters under each branch) represent bumble bee species assemblages (richness and abundance) and the environmental variable values associated with the study sites.

The multivariate regression tree for the northern plains geographic region resulted in a four-leaf tree where branching was determined by precipitation of driest month and minimum temperature of coldest month (Error = 0.571, CV Error = 1.48, SE = 0.362). *Bombus pensylvanicus* was the most abundantly collected species. Bumble bee assemblages were most rich in terms of both richness and abundance at sites with lower precipitation of driest month (Fig. 7A), containing 119 bumble bees across five species. Meanwhile, sites with medium levels of precipitation of driest month and lower minimum temperature of coldest month had 24 bumble bees present across three species (Fig. 7B). Similarly, 36 bumble bees from three species were found at sites with higher levels of precipitation of driest month and lower minimum temperature of coldest month (Fig. 7C). Comparably, sites with higher levels of precipitation of driest month and higher minimum temperature of coldest month had 28 bumble bees present representing four species (Fig. 7D), with *B. pensylvanicus* being the dominant species.

**Figure 7.**
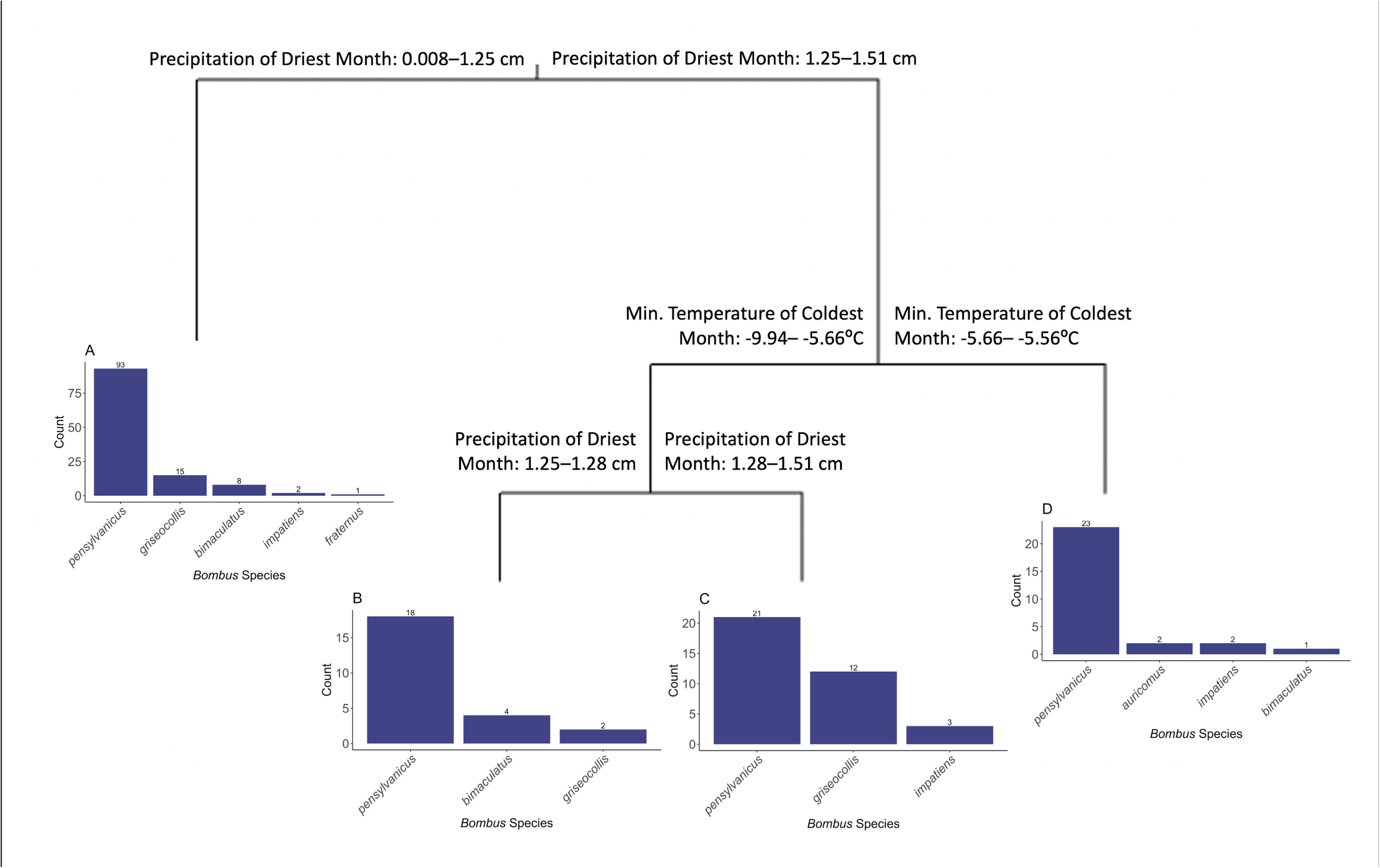
Multivariate regression tree for the northern plains geographic region, which resulted in branching at precipitation of driest month and minimum temperature of coldest month. The four leaves (indicated with letters under each branch) represent bumble bee species assemblages (richness and abundance) and the environmental variable values associated with the study sites.

The multivariate regression tree for the mountain geographic region resulted in a five-leaf tree where branching was determined by precipitation of wettest month, contiguity, bumble bee attractive crops, and developed land (Error = 0.438, CV Error = 0.797, SE = 0.151). *Bombus fervidus* was consistently the most abundant species across all leaves. Sites with lower precipitation of wettest month had the highest species richness, with 307 bumble bees across eleven species (Fig. 8A). Meanwhile, sites with higher precipitation of wettest month, lower contiguity, and less bumble bee attractive crops had 176 bumble bees present representing 10 species (Fig. 8B). Similarly, sites with higher precipitation of wettest month, lower contiguity, and more bumble bee attractive crops had 289 bumble bees present across 10 species (Fig. 8C). Moreover, 333 bumble bees from 9 species were present at sites with higher precipitation of wettest month, higher contiguity, and less developed land (Fig. 8D). Likewise, 452 bumble bees from 9 species were present at sites with higher precipitation of wettest month, higher contiguity, and more developed land (Fig. 8E).

**Figure 8.**
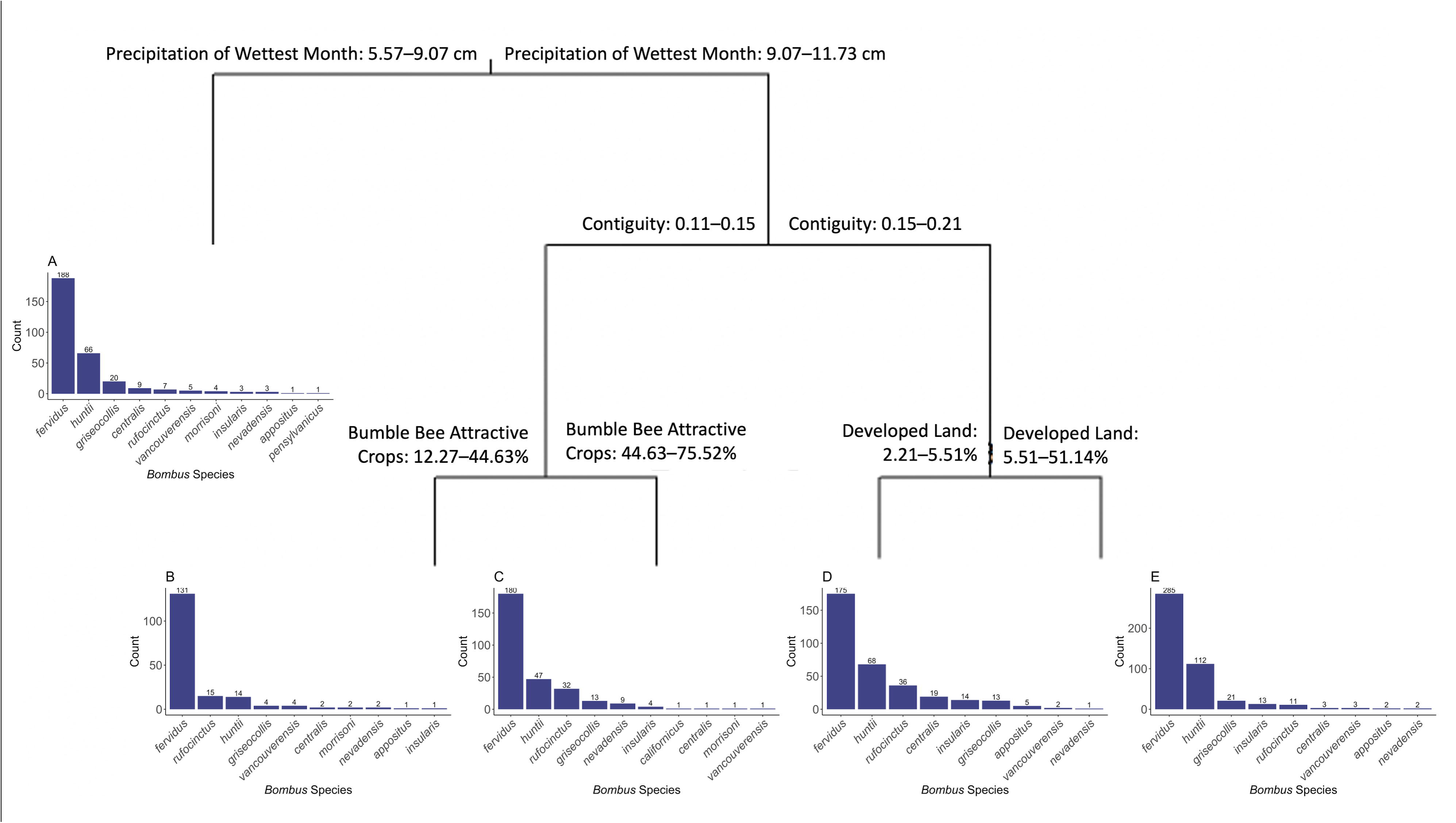
Multivariate regression tree for the mountain geographic region, which resulted in branching at precipitation of wettest month, contiguity, bumble bee attractive crops, and developed land. The five leaves (indicated with letters under each branch) represent bumble bee species assemblages (richness and abundance) and the environmental variable values associated with the study sites.

## 4. DISCUSSION

Overall, in this study we tested bumble bee responses to climate and land use by modeling 1) occupancy (presence/absence); 2) species richness; 3) species-specific habitat requirements; and assemblage-level responses and how those responses vary across geographic regions. Similar to previous studies, we found that temperature, precipitation, and land use significantly drive bumble bee assemblages (Naeem et al., 2019). Overall, climate and land use combine to drive bumble bee occupancy and assemblages, but how those processes operate is idiosyncratic and spatially contingent across regions.

### 4.1. Bumble Bee Occupancy and Richness Characterized by Climate and Landscape

We found that maximum temperature of warmest month, precipitation of wettest month, elevation, bumble bee attractive crops, bumble bee unattractive crops, interspersion and juxtaposition, and contiguity were the most important factors for predicting bumble bees occupancy and species richness. Increased prevalence of bumble bee attractive crops increased the odds of bumble bee presence, while also positively impacting the number of individual species found within the habitat. Meanwhile, increased prevalence of bumble bee unattractive crops and maximum temperature of warmest month increased the odds of bumble bee absence. Unlike our findings in the logistic regression that documented the absence of bees within increased maximum temperature of warmest month, the generalized additive mixed model provided more information on species richness trends that were not solely negative. Species richness increased with increasing maximum temperature of warmest month until 29°C before a marked decline as temperatures rose above 33°C. Therefore, while bumble bees were found at sites with temperatures between 33–36°C, this range may represent a temperature threshold for some species. From a management perspective, these findings highlight actions land managers can actively employ to increase the presence and richness of bumble bees within and around their farms. Planting bumble bee attractive crops such as tomatoes, peppers, cucumbers, berries, melons, squash, and soybeans surrounding less-bee friendly row crop fields (e.g., corn, small grains) can provide nutritive pollen and nectar for bumble bees, thereby increasing their presence and richness (USDA, 2017). Further, planting bumble bee attractive crop trees (i.e., peaches, cherries, apples, oranges, nectarines, plums, and apricots) for shade cover can help to provide microclimates to help offset the negative association with high maximum temperature of warmest month, while also increasing the presence of pollen and nectar within the environment for bumble bees. Further, while not significant in the logistical regression, precipitation of wettest month, elevation, and contiguity were again identified as being of importance to bumble bee species richness within the generalized additive model. Like the logistic regression, species richness declined with increased precipitation of wettest month, which may be a result of reduced foraging during these periods due to increased rainfall (Peat and Goulson, 2005; Sanderson et al., 2015). Additionally, as elevation increased there was a respective linear increase in species richness, which is likely a byproduct of geographic region (east vs. west) and the number of species found within the respective region. Increased contiguity led to a slight increase in species richness before declining as contiguity rose above 0.2. With this, it is important to note that contiguity was low throughout our study sites, meaning the landscapes were comprised of several different land cover types (Hesselbarth et al., 2019), which could increase the amount of floral and nesting resources available to bumble bees within the surrounding area.

### 4.2. Bumble Bee Habitat Associations

Species-environment associations identified that geographic region played a large role in species-specific habitat requirements, with distinct groupings in the east and west. In the east, species were associated with lower elevations along with high minimum temperatures of coldest month, increased precipitation of wettest month, and increased forest land surrounding the agricultural fields. These associations make sense because the east typically has lower elevations, warmer winters, and more precipitation compared to the west. It is important here to note that the west is solely represented by Utah, an intermountain state characterized by high elevation irrigated agricultural valleys, in this dataset. Therefore, results would likely be different if other western states were included in this dataset. In the west, Utah bees were associated with higher elevations, higher levels of attractive and unattractive crops, high maximum temperatures of warmest month, and increased interspersion and juxtaposition. Again, from a geographic standpoint these associations make sense because the west has higher elevations and warmer and drier summers than many of the eastern states, suggesting these bumble bees have adapted to these conditions. Further, the association of Utah bumble bee species with higher values of interspersion and juxtaposition suggests that these bumble bees are more abundant with landscapes that have a mix of land cover types that are well dispersed (less contiguous) within the surrounding area. One species that emerged as being of particular interest is *B. griseocollis,* which was ubiquitous throughout the study areas, following its known distribution (Colla et al., 2011; Koch et al., 2012). This species may be more resilient to land cover and climate change as they are able to survive well throughout a range of habitat types (i.e., open farmland and fields, urban parks and gardens, and wetlands) and climates across the U.S. (Koch et al., 2012; Kingsolver et al., 2013; Williams et al., 2014). Overall, this model identified that geographic region impacted species-specific habitat requirements, emphasizing the importance of evaluating species on a regional scale.

### 4.3. Climate and Landscape Impact on Bumble Bee Assemblages by Geographic Region

In the corn belt/Appalachian/northeast, sites were grouped by developed land, minimum temperature of coldest month, and maximum temperature of warmest month. Sites with higher proportions of developed land within the surrounding area supported rich bumble bee assemblages in terms of both richness and abundance. This may be a result of roadside strips, residential gardens, community gardens, and city parks having greater abundance of floral resources than the commodity crop fields surveyed in this study (Martins et al., 2017; Bennett and Lovell, 2019; Kelemen and Rehan, 2020; Wenzel et al., 2020). Further, the bumble bee species found in the sites with more developed land within the surrounding area were relatively similar, so minimum temperature of coldest month and maximum temperature of warmest month did not play a large role in impacting species richness at these sites. However, high and low values of minimum temperature of coldest month and maximum temperature of warmest month did impact the overall abundance of each species found at these sites. For example, at the temperature extremes (lower minimum temperature of coldest month and higher maximum temperature of warmest month) species were collected at lower abundances. Regardless, this finding identifies that within sites with a higher proportion of developed land in the surrounding area, species are found throughout a range of temperatures. Meanwhile, sites with low proportions of developed land in the surrounding area supported low bumble bee assemblages that were largely dominated by *B. bimaculatus.* Overall, these findings suggest that developed land surrounding agricultural fields in the corn belt/Appalachian/northeast region of the USA supports diverse bumble bee assemblages. However, further research is needed to determine what is driving this relationship, be it increased floral resource diversity, nesting sites, or reduced exposure to pesticides that would typically be found in agriculturally intensified settings (Goulson et al., 2015; Rundlöf et al., 2015; Janousek et al. 2022).

In the southeast, sites were grouped by forest land, precipitation of wettest month, and maximum temperature of warmest month. Sites with lower forest land, higher precipitation of wettest month, and lower and higher values of maximum temperature of warmest month effectively contained no bumble bees. This was likely due to the climatic extremes at these sites, with temperatures ranging from 32.29–34.17°C and precipitation ranging from 26.36–28.51 cm. While this temperature range is well below critical thermal maximas for many species (Oyen et al., 2016; Christman et al., 2022b), this is still a high temperature for cold-adapted bumble bees to sustain if held constant over long periods of time. Further, the high levels of precipitation at these sites likely prevents foraging in these fields and may even result in the flooding of nearby colonies, resulting in a lack of bumble bees at these sites (Peat and Goulson, 2005; Sanderson et al., 2015; Goulson et al. 2018). Interestingly, we found that sites with lower forest land and lower precipitation of wettest month supported bumble bee assemblages comprised of *B. pensylvanicus, B. impatiens, B. bimaculatus,* and *B. auricomus*. This finding may support the concept that precipitation instead of temperature plays a larger role in determining whether bumble bees will be present or absent at sites in Florida and South Carolina; however, further research is needed. Additionally, bumble bees were more species rich at sites with higher forest land in the surrounding area, which may be due to the increased availability of nesting sites within these habitats (Betts et al., 2019; Pfeiffer et al., 2019). Overall, these findings suggest that forest land increases bumble bee species richness and abundance, while extreme temperatures and precipitation can reduce the prevalence of bumble bees, emphasizing the importance of maintaining diverse forest habitats around agricultural fields on top of establishing climate mitigation techniques to reduce the magnitude of bumble bee declines (Prestele et al., 2021)

In the northern plains region, sites were grouped by precipitation of driest month and minimum temperature of coldest month. Bumble bee assemblages were richer in terms of richness and abundance at sites with lower precipitation of driest month. However, it is important to note that all sites had low precipitation of driest month (precipitation of driest month < 1.51), suggesting bumble bees in Kansas are adapted to drier environments.

In the mountain region, sites were grouped by precipitation of wettest month, contiguity, bumble bee attractive crops, and developed land. Sites with higher precipitation of wettest month, less contiguity, and the presence of bumble bee attractive crops supported species rich communities dominated by *B. fervidus.* Meanwhile, sites with higher precipitation of wettest month, more contiguity, and the presence of developed land supported less species rich communities with more individuals overall. Throughout the study sites, contiguity was low (0.11–0.21) meaning the sites were comprised of several different land cover types (Hesselbarth et al., 2019). Contiguity with the combination of bumble bee attractive crops and developed land is of particular interest as this suggests the habitats surrounding the surveyed agricultural fields were dominated by bumble bee attractive crops and developed land, which again likely supports bumble bee assemblages through an influx of more diverse floral resources. Overall, there was a great deal of species overlap between groupings. As such, bumble bees in Utah agroecosystems are likely supported by a range of climate, landscape composition, and landscape configuration variables. However, it is important to remember that all these species were collected in agricultural fields, which influences the composition of species collected within these surveys. Regardless, maintaining diverse habitats around agricultural fields can increase the richness and abundance of bumble bees.

### 4.4. Limitations

Although this study has some limitation inherent to its design (i.e., state-level differences in sample size, collection dates and period, target pests, and agriculturally focused trapping sites), analyzing trap bycatch reduces cost associated by allowing more efficient use of time and resources and utilizes data that would otherwise be discarded (Spears et al., 2021). Further, this study’s substantial spatial coverage and high number of replicates within and across years resulted in a large data set that enriches our knowledge of bumble bee assemblages across geographic space and time (Kohler et al., 2020). Additionally, the low proportion of singletons in this data set indicates a strong sampling regime (Williams et al., 2001; Kohler et al., 2020). Finally, the inclusion of climatic and landscape composition and configuration variables into one model introduces sources of uncertainty but yields more realistic results about their cumulative effects on *Bombus* species assemblages (Conlisk et al., 2013; Louca et al., 2015).

### 4.5. Broad-Scale Management Implications

This study further emphasizes the importance of conducting broad scale surveys. From this study, we identified that *B. pensylvanicus* was one of the most abundant species collected. This is of particular interest from a conservation perspective because its numbers have drastically declined since 2000 (∼89%) to the point that *B. pensylvanicus* is being evaluated to determine if it warrants federal protection under the U.S. Fish and Wildlife Service Endangered Species Act (USFWS, 2021). While the finding that *B. pensylvanicus* was one of our most abundant species may be due to their populations increasing, it could also be a result of this species being more attracted to pest monitoring traps. Further research is needed to determine which, if either, are true since both have significant management implications. If the former is true, *B. pensylvanicus* populations may be rebounding, so listing under the Endangered Species Act may not be warranted. If the latter is true, having an influx of *B. pensylvanicus* captured passively as bycatch could lead to declining populations. Regardless, this finding highlights the need for interagency cooperation and research collaborations to understand impacts of pest monitoring traps on bumble bee species and to develop and implement recovery plans and protective regulations.

While research is underway to understand the broader impacts of pest monitoring traps on bumble bees at a national scale (Spears and Ramirez, 2015; Spears et al., 2016, 2021; Christman et al., 2022a), science policy is needed to make actionable change. With informed science policy, innovative management practices and mandates can be employed that meet multiple federal agency’s goals, while also preventing the least amount of harm to agriculture, natural resources, and imperiled species. Further, given the diversity of insects captured within pest monitoring traps, improving policies, and increasing interagency collaboration could protect a range of insect and plant species, ecosystem services, and habitats.

### 4.6. Conclusions

Overall, results from this study contribute to a better understanding of climate and landscape factors affecting bumble bees and their habitats throughout the USA. Climate and land use combine to drive bumble bee occupancy and assemblages, but how those processes operate is idiosyncratic and spatially contingent across regions. Detailed knowledge of species-specific relationships with climate and landscape variables across a geographic range is invaluable to improve targeted conservation and land management strategies to mitigate the effects of ongoing environmental changes.

## ACKNOWLEDGEMENTS

We thank Harold Ikerd for his assistance with data management; Soli Velez, Kami Lay, and Anna Fabiszak for their assistance with processing insects; Todd Gilligan, Brian Christman, Zachary Schumm, and the anonymous reviewers for invaluable comments that improved this manuscript; and the following state cooperators for providing bycatch samples: Larry Bledsoe, Bradley Danner, Eric Day, David Gianino, Mathew Howle, Janet Lensing, Laura Miller, Laurinda Romonda, and Gaye Williams. This work was made possible, in part by Cooperative Agreements AP18PPQFO000C100, AP19PPQFO000C269, AP19PPQS&T00C056, AP20PPQFO000C074, AP20PPQS&T00C065, and AP21PPQS&T00C056 from the United States Department of Agriculture’s Animal and Plant Health Inspection Service (APHIS) and National Science Foundation Grant No. 1633756. Work may not necessarily express APHIS’ views.

## AUTHOR CONTRIBUTION

Conceptualization: MEC, EKB, LRS, WPD, and RAR. Data curation: MEC. Formal analysis: MEC, EKB, and WPD. Funding acquisition: LRS and RAR. Investigation: MEC. Methodology: MEC, EKB, LRS, WPD, and RAR. Project administration: LRS and RAR. Resources: LRS and RAR. Supervision: EKB, LRS, WPD, JPS, and RAR. Writing – original draft: MEC. Writing – review and editing: EKB, LRS, WPD, JPS, and RAR.

## DATA AVAILABILITY STATEMENT

Data and code supporting the findings of this study are available on Zenodo at https://doi.org/10.5281/zenodo.6363812.

